# Type I IFN promotes pathogenic inflammatory monocyte maturation during H5N1 infection

**DOI:** 10.1101/2022.01.10.475751

**Authors:** Slim Fourati, David Jimenez-Morales, Judd Hultquist, Max W. Chang, Christopher Benner, Nevan Krogan, Lars Pache, Sumit Chanda, Rafick-Pierre Sekaly, Adolfo García-Sastre, Melissa B. Uccellini

## Abstract

Ly6C^hi^ inflammatory monocytes show high IFN responses, and contribute to both protective and pathogenic functions following influenza virus infection. In order to understand the significance of IFN responses in this subset, we examined monocytes during infection with a lethal H5N1 virus that induces high levels of IFN and a low-pathogenicity H1N1 virus that induces low levels of IFN. We show that H5N1 infection results in early recruitment of high numbers of Ly6C^hi^ monocytes and induction of chemokines and *Ifnb1*. Using unbiased transcriptomic and proteomic approaches, we also find that monocytes are significantly enriched during H5N1 infection and are associated with chemokine and IFN signatures in mice, and with severity of symptoms after influenza virus infection in humans. Recruited Ly6C^hi^ monocytes subsequently become infected in the lung, produce IFN-β, and mature into FasL^+^ monocyte-derived cells (FasL^+^MCs) expressing dendritic cell markers. Both *Ccr2^-/-^* and *Fasl^gld^* mice are protected from lethal infection, indicating that monocytes contribute to pathogenesis. Global loss of type I and type III IFN signaling in *Stat2^-/-^* mice results in loss of monocyte recruitment, likely reflecting a requirement for IFN-dependent chemokine induction. Here we show that IFN is not directly required for monocyte recruitment on an IFN-sufficient background, but is required for maturation to FasL^+^MCs. Loss of IFN signaling skews to a Ly6C^lo^ phenotype associated with tissue repair, suggesting that IFN signaling in monocytes is a critical determinant of influenza virus pathogenesis.

## Introduction

Monocytes are hematopoietic cells that develop from myeloid progenitors in the bone marrow and traffic via the bloodstream to peripheral tissues. Chemokine (C-C motif) receptor 2 (CCR2) is required for their release from the bone marrow into the blood, in a process dependent on both chemokine (C-C motif) ligand 2 (CCL2) and CCL7. Recruitment of monocytes from the blood to the tissues is independent of CCR2 and likely involves other chemokines (1–3). Monocytes are defined by expression of CD115 and CD11b, and are divided into two subsets termed “inflammatory” and “patrolling” monocytes. Inflammatory monocytes express high levels of Ly6C and CCR2 in mice, and are CD14^+^CCR2^+^CD16^-^ in humans. Patrolling monocytes express low levels of Ly6C, no CCR2 and are GFP^hi^ in CX3CR1^GFP^ mice, and are CD14^lo^CX3CR1^hi^CD16^+^ in humans. Inflammatory signals lead to recruitment of Ly6C^hi^ monocytes to sites of infection, where they differentiate into various tissue macrophage and dendritic cell populations that aid in infection clearance. Ly6C^hi^ monocytes can give rise to Ly6C^lo^ monocytes (4–7), although development independent of Ly6C^hi^ monocytes is also supported by some studies (8, 9). The Ly6C^lo^ subset has been associated with wound healing and tissue remodeling following injury or infection (10–12).

During influenza virus infection, monocytes have been shown to contribute to both protective functions, and tissue damage when recruited in excess numbers. Monocyte populations that have differentiated into effector cells in the lung have been termed various things during influenza virus infection – including monocyte DCs, TNF-α/iNOS-producing DCs, inflammatory DCs, exudate macrophages, and inflammatory monocyte-macrophages (13–15). Currently there is no naming consensus for these populations, which has led to confusion in the literature. Here we follow the naming convention proposed by Guilliams (16) and refer to monocyte-derived cells differentiated in the influenza-infected lung as ‘FasL^+^MCs’ to denote that they are monocyte-derived cells (MCs) with the functional characteristic of FasL expression, in addition to DC markers including CD11c and MHC II. Importantly, they express CCR2, CD115 and CD64, and not CD24 indicating monocyte rather than dendritic cell origin. The populations described in different studies likely represent similar, but heterogeneous differentiated MC populations. Monocyte accumulation in the lung has also been described in many infections including SARS-CoV-1 and SARS-CoV-2 (17–19). Influenza-infected *Ccr2^-/-^* mice fail to recruit monocytes, and show delayed viral clearance, decreased T cell accumulation in the draining lymph node, and decreased CD8^+^ T cell priming (20, 21) supporting protective roles for monocytes. However, CCR2 deficiency protects from influenza-induced mortality in most studies (20, 22, 23), correlating with decreased lung injury – indicating that in many cases monocytes acquire pathogenic phenotypes and contribute to immune-mediated tissue damage.

Type I and type III IFN are critical for restricting viral replication and systemic dissemination, as evidenced by the high susceptibility of *Ifnar1^-/-^* mice to a number of viruses (24). However, *Ifnar1^-/-^* mice infected with influenza virus show increased or decreased susceptibility to infection depending on the experimental conditions (25), making the precise role of IFN during influenza virus infection *in vivo* more difficult to define. This likely reflects the ability of IFN to act in many different ways on a wide-range of cell types, and to induce the expression of ISGs, which orchestrate later immune cell infiltration. Factors that influence the balance between the protective and pathogenic functions of IFN include the background of the mouse strain, the presence or absence of functional Mx1 expression, the viral strain, and redundancy between the functions of type I and type III IFN. Highly susceptible 129 and DBA/1 strains produce high levels of IFN, which subsequently leads to immune-mediated tissue damage (26). Therefore lack of IFN-signaling during influenza virus infection in these strains is protective. Similar differences in pathogenesis in *Ifnar1^-/-^* mice have been reported for SARS-CoV-1, with IFN mediating detrimental effects in highly susceptible BALB/c mice (17), but not C57BL/6 or 129 mice (27, 28). Most inbred mouse strains lack expression of Mx1 (29), however on an Mx1-sufficient background IFN is protective through induction of Mx1 expression and potent restriction of influenza virus replication. Importantly, the increased susceptibility phenotype is only strongly evident in strains lacking both type I and type III IFN signaling due to redundancy in Mx1 induction (30). Certain viral strains have the ability replicate outside of the lung due to ability of the hemagglutinin protein to be cleaved independently of trypsin-like proteases. In these strains, lack of type I IFN signaling is detrimental and leads dissemination of virus outside of the lung (31–34). Given the many different phenotypes of influenza-infected *Ifnar1^-/-^* mice, understanding the function of IFN on specific cell types during influenza virus infection will be important for understanding pathogenesis.

Using an ISRE-dependent reporter mouse, we previously showed that Ly6C^hi^ monocytes have high IFN responses following influenza virus infection (35), however the significance and specific function of IFN in this subset is not known. During influenza virus infection, monocytes have been shown to accumulate in a manner dependent on virus pathogenicity and type I IFN levels (22, 36, 37). In addition, in susceptible mouse strains, high IFN levels lead to excessive monocyte recruitment. IFN-dependent expression of TRAIL by monocytes subsequently leads to damage to the lung epithelium (26, 38). Here we investigated the functional significance of IFN signaling in Ly6C^hi^ monocytes. We show that IFN-signaling is not directly required for recruitment, but is critical for maturation to a pathogenic phenotype. In the absence of IFN signaling monocyte phenotypes are skewed to a Ly6C^lo^ phenotype associated with tissue repair, indicating that IFN signaling in monocytes is critical for determining the balance between the protective and pathogenic functions of type I IFN during influenza virus infection.

## Results

### Ly6C^hi^ monocytes are associated with H5N1 infection in mice, and severity of symptoms in humans

Using an ISRE-dependent reporter mouse for type I and type III IFN (*Mx1^gfp^*), we previously showed that Ly6C^hi^ monocytes display very high IFN responses in the lung following influenza virus infection (35). In order to understand the significance of IFN responses in this subset, we examined monocytes during infection with either an avian H5N1 strain of influenza virus (A/Vietnam/1203/04 HALo) or a lab-adapted H1N1 infection (A/PR/8/34). The HALo strain has been engineered to remove the hemagglutinin multibasic cleavage site, which limits systemic spread of the virus and allows use under BSL2 conditions, however the virus retains its other virulence determinants (39). At the doses examined, H5N1 induces an early IFN response that is sustained until the animals succumb to infection - while H1N1 induces a delayed IFN response, and the animals go on to survive infection (Fig 1A) (35). At day 2 post-infection, mice infected with H5N1 showed a large influx of Ly6C^hi^ monocytes (Fig 1B). In contrast, mice infected with H1N1 showed Ly6C^hi^ monocyte numbers similar to uninfected mice. We were unable to detect consistent viral titers at day 2 post-infection in H5N1 or H1N1 infected animals, however day 3 post-infection both viruses replicated to high titers in the lung, (Fig 1C) suggesting viral replication did not account for the large differences in monocyte infiltration and infection. Using complementary unbiased transcriptomic and proteomic approaches, we compared the lung tissue of mice infected with the same H5N1 strain to another mild H1N1 strain (A/California/04/09) at 12 hours, or 1, 2, 3 or 4 days post-infection. In order to identify immune cell subsets that may be differentially engaged by H5N1 and H1N1, we assessed the enrichment (i.e. overlap) between known transcriptomic (40) and proteomic (41) markers of immune cell subsets (T, B, NK, monocyte, mDC, pDC) among genes/proteins differentially expressed between the two strains. Monocytes were the only significantly enriched immune cell subset associated with H5N1 infection by transcriptomics (Fig 1D) and proteomics (Fig S1). Monocytic genes and proteins induced by H5N1 included the cell surface markers Cd14 and Fcgr1a as well as the interleukin Il15 and TLR4-ligand S100a9. In order to determine if the monocytes gene signature induced after H5N1 infection in lung of mice can also be detected in blood, we analyze an independent set of mice infected with H3N2 for which both lung and blood RNA-sequencing was performed [PMID:26413862]. The monocytes signatures was induced post-infection and reach it maximum induction 3 days following infection in both lung and blood.

**Figure 1.**
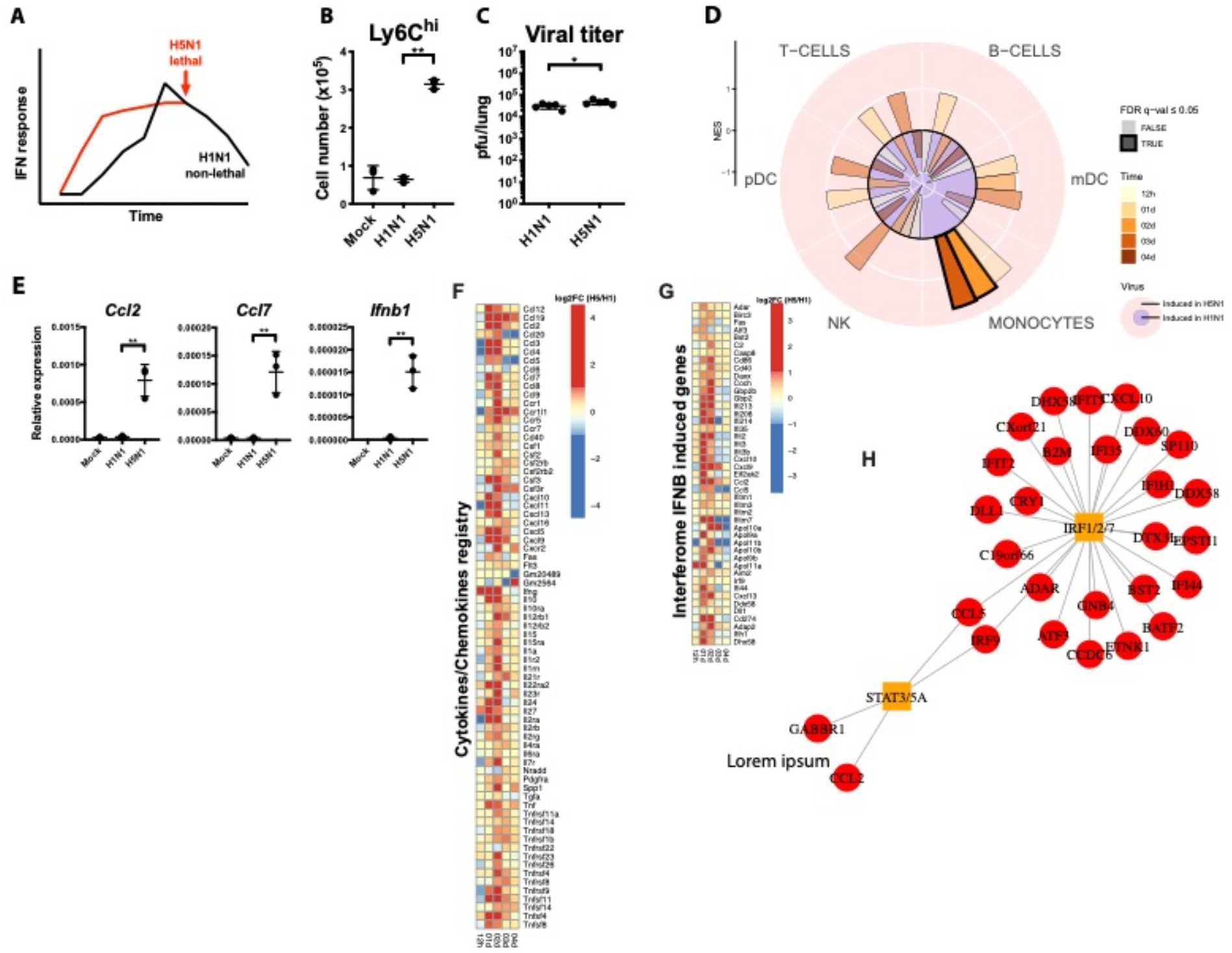
Monocytes are associated with H5N1 infection. **(A)** IFN response and pathogenesis of H1N1 and H5N1 at the doses used in this study **(B)** Mice were infected with 10^2^ pfu of H1N1 or H5N1 and Ly6C^hi^ monocyte (Ly6C^hi^CD11b^+^Ly6G^-^) infiltration and was determined by flow cytometry at day 2 post-infection (n=3). **(C)** Mice were infected as in B and viral titer was determined by plaque assay at day 3 post-infection. Data are representative of 3 experiments for B and 2 experiments for C. *p<0.05**p<0.01 **(D)** Gene Set Enrichment analysis was performed to assess the enrichment of transcriptomic markers of immune cells (B, T, NK, Monocytes, mDC, pDC; (40) among genes differentially expressed between H5N1- and H1N1-infected mice in lung tissue. The radial plot presents the Normalized enrichment score (NES) of each set of markers (quadrants) for different timepoints investigated after infection (bars). A NES > 0 correspond to enrichment of cell markers among genes induced in H5N1 compared to H1N1 while a NES < 0 corresponds to the enrichment of cell markers among genes repressed in H5N1 compared to H1N1. A permutation test was performed to assess the significance of each enrichment; opaque bars correspond to enrichment with a false discovery rate (FDR) below 0.05. **(E)** Mice were infected with 10^2^ pfu of H1N1 or H5N1 and Ly6C^hi^ monocyte (Ly6C^hi^CD11b^+^Ly6G^-^) infiltration and chemokine induction was determined by qPCR at day 2 post-infection (n=3). Data are representative of 3 experiments. **p<0.01 **(F)** Heatmap showing the log2 fold-change (log2FC) of genes coding for cytokines/chemokines between H5N1- and H1N1-infected mice in lung tissue. A blue-white-red color gradient depicts the gene the most repressed to the gene the most induced in H5N1-infected mice compared to H1N1-infected mice. **(G)** Gene Set Enrichment Analysis (GSEA) was performed to assess the enrichment (i.e. overlap) of interferon-stimulated genes (IFN-a, β, λ, and γ induced genes; (80)) among genes differentially expressed between H5N1- and H1N1-infected mice in lung tissue. This analysis revealed a significant enrichment of IFN-β-induced genes in the lungs of H5N1-infected mice on day 2. The heatmap shows the log2FC of genes contributing to the IFN-β-induced gene enrichment in H5N1-infected mice compared to H1N1-infected mice. **(H)** GSEA was used to identify transcription factors (TF) upstream of IFN-β-induced genes in the lungs of H5N1-infected mice on day 2. In the network, TF (scare) and their targets (circle) are linked (edge) if there is a putative binding sites in the 2000 bases around the TSS of the target genes or that the genes was differentially regulated in TF-overexpression or knock-out experiments.

Subsequently, we investigated the use of the monocytes gene signature induced after H5N1 as being associated with influenza disease severity in human in a large meta-analysis of blood transcriptomic datasets [PMID: 30356117]. Fig 2). Similarly to the mice lung, the monocytes gene signature was induced day 2 to day 4 after infection in participants that developed severe symptoms. Genes in the monocyte signature that were most significantly associated with severity in human include the chemokine coding genes CXCL10 and CCL2. These subjects showed heightened expression of the TNF/NFκb confirming the pro-inflammatory environment triggered by the infection. Markers of monocytes (CD86, CCR5) and inflammasome activation (CASP1, NLRP3) required to activate pro-IL-1β and IL-18 were also associated with disease severity. Counterintuitively, this gene signature also included increased expression of the immunomodulatory cytokine IL-10 well known to be expressed by monocytes most probably as a negative feedback loop to overcome the heightened inflammatory environment triggered by infection. Similarly to H5N1 infected mice lung severe symptoms. The signature resolved by day 7 in all infected subjects. Altogether, these results highlight the pro-inflammatory environment triggered by H1N1 infection in human subjects which is largely attributed to the activation of cells of the monocyte lineage. The conservation of these gene signatures between mice and human infected with the flu virus and the association with disease severity suggests that monocyte activation is a hallmark feature of flu infection pathogenesis.

**Figure 2.**
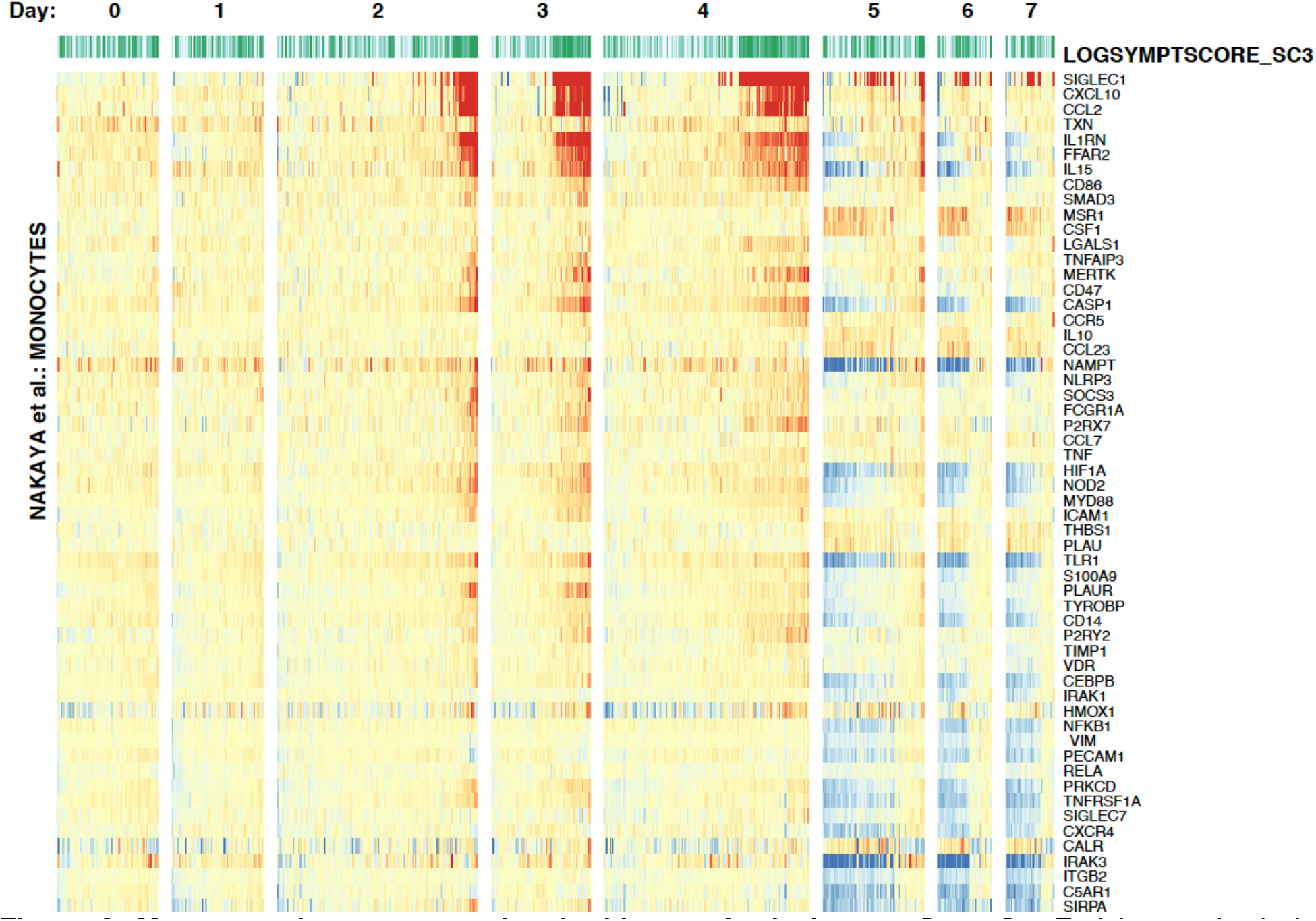
Monocyte signature associated with severity in human. Gene Set Enrichment Analysis (GSEA) was performed to assess the enrichment (i.e. overlap) of transcriptomic markers of immune cells (B, T, NK, Monocytes, mDC, pDC) among genes expressed in blood of healthy adults challenged with H1N1 and correlated with severity of symptoms (LOGSYMTOMSCORE; (79)). A significant enrichment of monocyte signature was observed on days 2, 3 and 4. The heatmap shows the expression of transcriptomic markers of monocytes contributing to this enrichment. A blue-white-red color gradient represents down-regulation to up-regulation of the gene-expression scaled across samples (z-score).

### H5N1 infection results in early chemokine and Ifnb1 production

Ly6C^hi^ monocytes produce the CCR2 ligands CCL2 and CCL7 in an IFN-dependent manner, which further amplifies their recruitment (17, 22). Consistent with this, animals infected with H5N1 showed robust induction of *Ccl2* and *Ccl7*, while mice infected with H1N1 did not show induction of these chemokines (Fig 1E). Pathway enrichment analysis of the genes and proteins differentially expressed in H5N1 infected mice compared to H1N1 infected mice confirmed that *Ccl2* and *Ccl7*, as well as a number of other chemokines and cytokines are induced at early timepoints post-infection (Fig 1F).

Transcriptomic analysis also revealed a strong IFN signature at early time points in H5N1 infected mice. In addition, mice infected with H5N1, but not H1N1 showed robust induction of *Ifnb1* (Fig 1G), consistent with a report showing *Ifnb1* message in sorted Gr1^+^CD11b^+^ cells (22). compared to H1N1 infected mice (Fig 2C). Interferon-stimulated genes induced in H5N1 infected mice include the viral sensors Dhx58 and Ifih1 and the transcription factor Irf9 (Fig 1H). This is consistent with the reported ability of H5N1 strains to induce a “cytokine storm (42, 43).

### Ly6C^hi^ monocytes become infected and produce IFN-β during H5N1 infection

We next examined the phenotype of Ly6C^hi^ monocytes following recruitment to the lung. At day 2 post-infection, H5N1-infected animals showed high numbers of Ly6C^hi^ monocytes that were infected as indicated by surface expression of influenza M2 protein, consistent with other reports showing that monocytes can become infected with influenza virus (22, 44)(28555541). By day 2 post-infection, H5N1-infected animals showed high frequencies of GFP^hi^ cells in H5N1-infected *Mx1^gfp^* mice; H1N1 infected mice showed significantly lower numbers (10 fold) of GFP^hi^ cells than H5N1 infected mice (Fig 3A and 3B). Kinetic analysis showed that H5N1 infected mice showed significancy higher frequencies of We previously observed a lack of *Ifnb1* induction in *Stat2^-/-^* mice (35), which was surprising given that *Ifnb1* is induced independently of IFN signaling. Lack of *Ifnb1* induction in both H5N1-infected *Stat2^-/-^* mice and H1N1 infected WT mice (Fig 2A) correlated with lack of infiltration of Ly6C^hi^ monocytes, suggesting that these cells might be a primary source of *Ifnb1* during infection. In order to determine if Ly6C^hi^ monocytes produce IFN-β, we infected *Ifnb^mob^* mice, which express YFP from an internal ribosome entry site in the *Ifnb1* locus, with H5N1 and monitored cells in the lung for YFP expression. We were able to detect YFP expression in ~18% of Ly6C^hi^ monocytes, but not in other cell types including xxxx (Fig 2C), suggesting that Ly6C^hi^ monocytes are indeed the main source of IFN-β during influenza virus infection. We were unable to detect YFP expression at various time points post-infection with H1N1 (not shown), indicating that IFN-β induction during H1N1 infection was below the limit of detection of the reporter gene. Ly6C^hi^ monocytes were also GFP^hi^ in H5N1-infected *Mx1^gfp^* mice, indicating they both produced, and responded to type I IFN (Fig 3A and D). Overall the data suggest that during H5N1 infection, Ly6C^hi^ monocytes recruited to the lung become infected and both produce and respond to type I IFN to levels higher than other cell types present in the lung. High levels of type I IFN production by Ly6C^hi^ monocytes in the lung likely contribute to increased infiltration of additional Ly6C^hi^ monocytes by promoting expression of recruiting chemokines, contributing to a pathological feedback loop.

**Figure 3.**
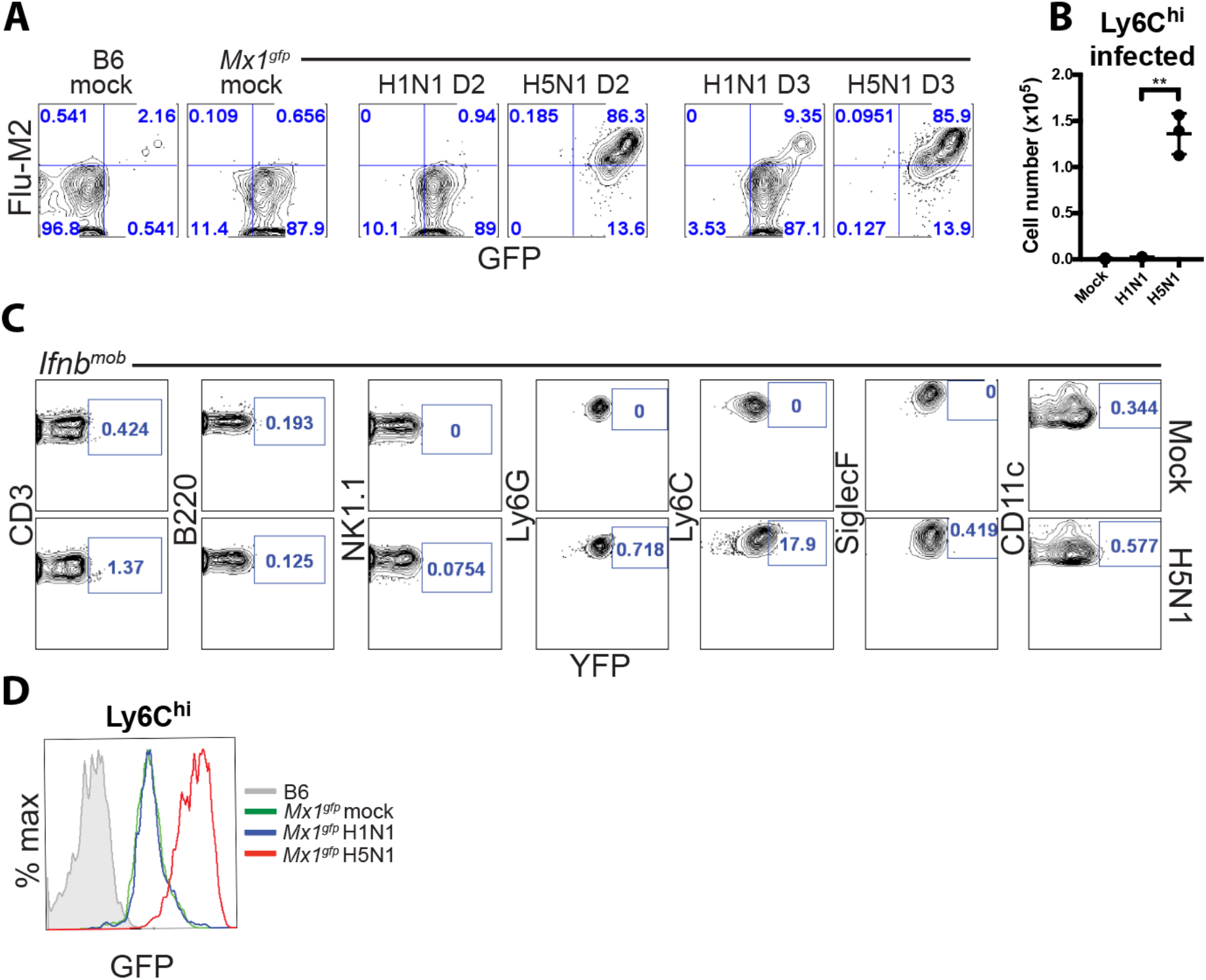
Monocytes become infected and produce IFN-β and ISGs. **(A)** *Mx1^gfp^* mice were infected with 10^2^ pfu of H1N1 or H5N1 and surface Flu-M2 expression on Ly6C^hi^ monocytes was determined by flow cytometry at day 2 and 3 post-infection. **(B)** Quantification of A (n=3). Data are representative of 3 experiments. **p<0.01. **(C)** *Ifnb^mob^* mice were infected with 10^2^ pfu of H5N1 and YFP expression was measured by flow cytometry at day 3 post-infection (n=2). Data are representative of 3 experiments. **(D)** *Mx1^gfp^* mice were infected with 10^2^ pfu of H1N1 or H5N1 and GFP expression was measured in Ly6C^hi^ monocytes. Data are representative of 3 experiments.

### Ly6C^hi^ mature into FasL^+^ MCs during H5N1 infection

Following recruitment to sites of infection, Ly6C^hi^ monocytes mature into various effector cells with different functional characteristics. When we further examined the phenotype of Ly6C^hi^ monocytes from H5N1-infected mice, we found they expressed higher levels of Ly6C, and upregulated CD11c and MHC Class II (Fig 4A). Influenza infection of 129 mice leads to high levels of IFN production and expression of TRAIL on Ly6C^hi^ monocytes, leading to subsequent DR5-dependent tissue damage (23, 26). We did not detect TRAIL expression on Ly6C^hi^ monocytes during H1N1 or H5N1 infection (not shown). FasL expression has also been reported to be upregulated during influenza virus infection (45, 46). When we examined FasL expression on Ly6C^hi^ monocytes, we found FasL was upregulated on Ly6C^hi^ monocytes at day 2 post-infection with H5N1 but not H1N1 (Fig 4A). We observed constitutive Fas expression on both hematopoietic and non-hematopoietic cells (not shown) as reported (45). The data suggest H5N1 infection leads to Ly6C^hi^ monocyte infection, IFN-β production, and maturation to a FasL^+^MC phenotype.

**Figure 4.**
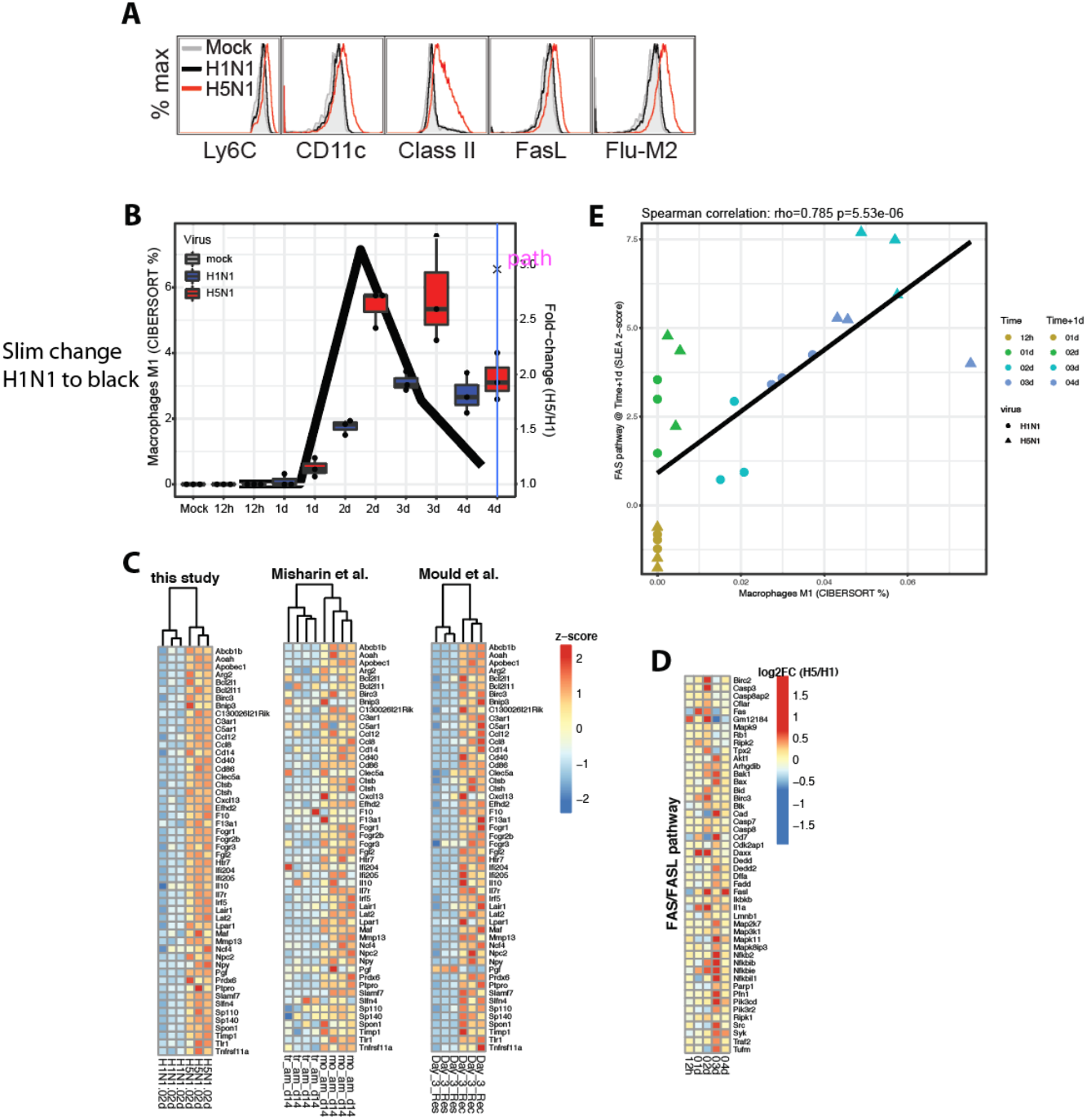
Monocytes mature into M1/FasL+MC phenotype during severe infection. **(A)** Mice were infected with 10^2^ pfu of H1N1 or H5N1 and expression of the indicated markers on Ly6C^hi^ monocytes was determined at day 2 post-infection (n=3). Data are representative of 3 experiments. **p<0.01 (**B**) Cibersort (81) was used to infer cell frequencies based on RNA-Seq data. Boxplot of inferred Macrophage M1 (IL-12^high^, IL-23^high^, IL-10^lo^^w^ phenotype, polarized by LPS) frequencies in lung of H5N1-, H1N1- and mock-infected mice at different timepoints following infection. (**C**) Transcriptomic expression of genes induced in monocyte-derived macrophages compared to tissue resident macrophages in the lung of H5N1- and H1N1-infected mice. GSEA enrichment analysis was used to test for enrichment of transcriptomic markers of monocyte-derived macrophages and of tissue resident macrophages (82, 83). A significant enrichment of monocyte-derived macrophage markers among genes induced in H5N1- vs. H1N1-infected mice. Genes contributing to the enrichment (i.e. overlapping monocyte-derived macrophages and induced in H5N1 infected mice) are shown in the heatmaps. A blue-white-red color gradient represents down-regulation to up-regulation of the gene-expression scaled across samples (z-score). (**D**) Gene Set Enrichment Analysis (GSEA) was performed to assess the enrichment (i.e. overlap) of transcriptomic markers of cell death pathways (Apoptosis, Pyroptosis, Necrosis) among genes differentially expressed between H5N1- and H1N1-infected mice in lung tissue. This analysis revealed a significant enrichment of Fas-mediated cell death in the lung of H5N1-infected mice at day 3. The heatmap shows the log2FC of genes contributing to the Fas pathway enrichment in H5N1-infected mice compared to H1N1-infected mice. (**E**) Scatter plot showing the levels of Fas pathway (summarized using the SLEA z-score method) in the lung of H5N1-, H1N1- and mock-infected mice as a function of the inferred frequency of M1 macrophage (inferred from Cibersort) the preceding day. A Spearman correlation and t-test was used to statistically assess the correlation.

In order to determine if our transcriptomic data similarly supported the maturation of infiltrating monocytes, we used CIBERSORT a method that uses gene expression profiles to quantify the distribution of hematopoietic cell susbets based on gene expression profiles. This method infers 22 cell subsets – including monocytes, M0 (differentiated from monocytes), M1 (differentiated in LPS/IFN-γ), and M2 (differentiated in LPS/IFN-γ/IL-4) cells for the monocyte/macrophage lineage. Deconvoluting our transcriptional data using this method, we found an enrichment of “classically activated” M1 macrophages in H5N1-infected mice, which peaked at day 2 post-infection (Fig 4B). Importantly the “M1” classification denotes a proinflammatory phenotype, consistent with our flow cytometry findings, as opposed to the repair phenotype that defines the M2 subset. In order to determine if these M1 macrophages were derived from tissue-resident macrophages, or included mostly infiltrating monocyte-derived macrophages, we performed GSEA enrichment analysis to compare our transcriptomic signatures to known markers of tissue-resident vs. infiltrating macrophages. This analysis revealed the significant enrichment of monocyte-derived macrophage markers in the lungs of H5N1, compared to H1N1 infected mice supporting their origin from infiltrating cells (Fig 4C). Of note these recruited monocytes exhibited heightened expression of several markers of monocyte activation (Cd86, Cd40), markers of the complement pathways (C1), and chemokines including Ccl8 and Cxcl13 and as well Il10 and Arg2 two cytokines with anti-inflammatory activity reiterating that a negative feedback loop is put in place to counteract potential deleterious effect of the heightened pro-inflammatory response. Of note these cells express also several genes with anti-apoptotic activity (Bcl-2, Birc3) suggesting that they are resistant to cell death signals prevalent in his pro-inflammatory environment

In order to determine if FasL expression was associated with M1 macrophages, we assessed the correlation between the frequency of M1 macrophages post-influenza infection, and cell death-specific pathways. This analysis showed enrichment for genes in the FasL pathway (Fig 4D) and a positive correlation of the FasL pathway with the inferred frequency of M1 macrophages (Fig 4E and Fig S3). FasL pathway was induced after H5N1 infection and reached its peak induction at day 3 post-infection. FasL pathway follow the same kinetics of response to infection as the monocyte signature but delayed by 1 day; this suggest that FasL pathway induction follow the induction of the FasL pathway. To test that FasL pathway is induced more strongly in severe flu disease and follow the induction of the monocytes gene signature, we analysed a publically available transcriptomic dataset of mice infected by two strains of H3N2 (one strain with low virulence and one highly virulent strain) [PMID:21874528]. This analysis confirmed that FasL pathway induction occurs 1 day following induction of the monocytes gene signature (Fig S3A) and revealed that FasL pathway is more strongly induced (> 3 fold-difference) by a virulent influenza strain (Fig S3B). The data suggest H5N1 infection is associated with maturation of Ly6C^hi^ monocytes into a proinflammatory M1/FasL^+^MC phenotype

### Ly6C^hi^ monocytes contribute to lethality during H5N1 infection

During influenza virus infection, low-level monocyte recruitment is likely beneficial, while excessive recruitment can lead to lung injury. In order to determine whether monocyte recruitment was beneficial, or contributed to lethality during H5N1 infection, we infected WT and *Ccr2^-/-^* mice, which are unable to recruit monocytes from the bone marrow to the blood, with H5N1 and monitored survival. *Ccr2^-/-^* mice were completely protected following H5N1 infection (Fig 5A), suggesting they contribute to lung damage. In addition, *Fasl^gld^* mice were partially protected following H5N1 infection (Fig 5B), suggesting that FasL is also a contributing factor to lung damage during H5N1 infection.

**Figure 5.**
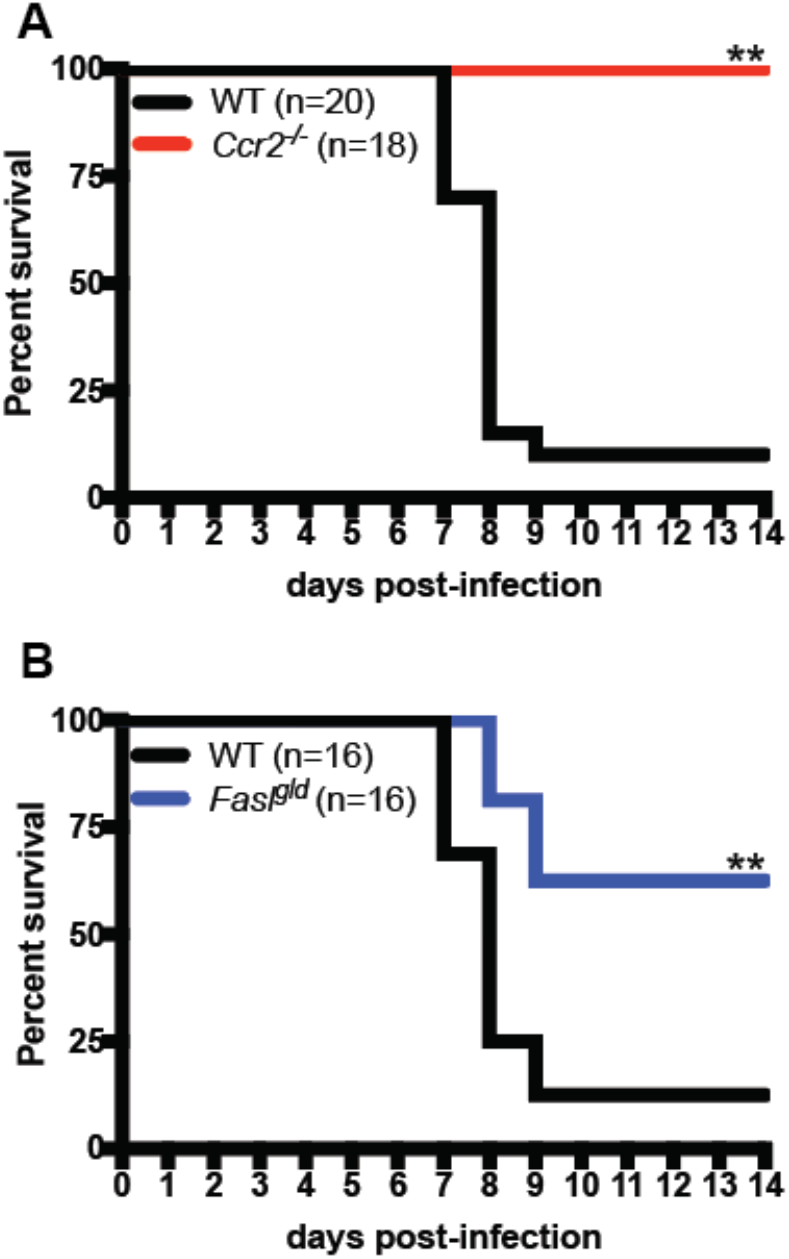
Monocytes contribute to lethality during severe infection. **(A)** WT and *Ccr2^-/-^* mice were infected with 10 pfu of H5N1 and survival was measured. **(B)** WT and *Fasl^gld^* mice were infected as in A and survival was measured. Data show combined results of 2 experiments with similar results. **p<0.01.

### IFN-signaling is not directly required for monocyte recruitment

Using *Stat2^-/-^* mice, we previously reported that Ly6C^hi^ monocyte recruitment is dependent on IFN signaling (35). Similar findings have also been reported in *Ifnar1^-/-^* mice (22, 36). Importantly, both *Ifnar1^-/-^* and *Stat2^-/-^* mice fail to induce ISGs including *Ccl2* and *Ccl7* (35, 36), which are required for the recruitment of Ly6C^hi^ monocytes from the bone marrow to the blood. In order to address if IFN-signaling was directly required for Ly6C^hi^ monocyte recruitment into the blood, we injected WT and *Stat2^-/-^* mice i.v. with CCL7 (47) and measured Ly6C^hi^ monocyte recruitment to the blood. Although *Stat2^-/-^* mice had lower steady-state levels of Ly6C^hi^ monocytes present in the blood, they were able to mobilize Ly6C^hi^ monocytes from the bone marrow to the blood to the same extent as WT mice (Fig 6A). CCL2 injection in the lung has been reported to recruit Ly6C^hi^ monocytes (48), however we were unable to reproduce these results. This is in agreement with work suggesting that CCL2 is required to recruit Ly6C^hi^ monocytes from the bone marrow to the blood, however it is not required for recruitment from the blood to the tissues, suggesting that other signals mediate tissue entry from the blood (1–3).

**Figure 6.**
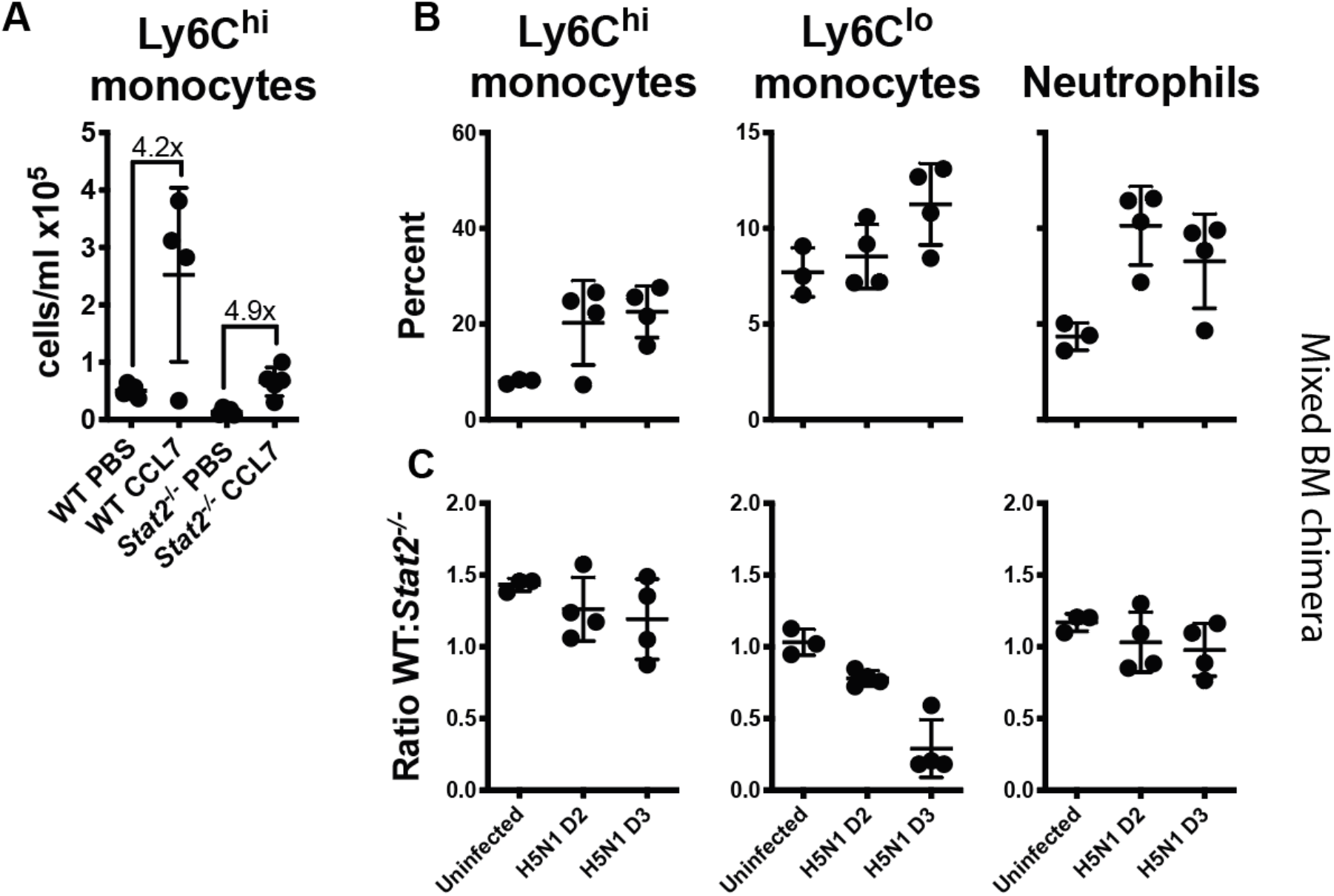
IFN-signaling is not directly required for monocyte recruitment. **(A)** WT and *Stat2^-/-^* mice were injected IV with CCL7 and Ly6C^hi^ monocytes in the blood were measured by flow cytometry at 30 min (n=4). **(B)** WT:*Stat2^-/-^→WT* 50:50 mixed bone marrow chimeras were infected with 10^2^ pfu of H5N1 and Ly6C^hi^ and Ly6C^lo^ monocyte, and neutrophil influx was measured by flow cytometry at day 2 and 3 post-infection (n=4). Data are representative of 2 experiments.

In a separate approach, we generated mixed bone marrow chimeras by reconstituting WT (CD45.1) mice with a 50:50 mix of WT (CD45.1) and *Stat2^-/-^* (CD45.2) bone marrow. In these mice, IFN signaling in both WT stromal cells and WT hematopoietic cells can induce CCL2 and CCL7 production. Following infection, mixed bone marrow chimeras show Ly6C^hi^ monocyte as well as neutrophil recruitment to the lung following H5N1 infection (Fig 6B). Approximately equal ratios of WT and *Stat2^-/-^* Ly6C^hi^ monocytes are present at day 2 and day 3 post-infection (Fig 6C). This suggests that IFN signaling in monocytes is not directly required for the recruitment of Ly6C^hi^ monocytes to the lung, but likely is required to induce chemokine production that subsequently leads to their recruitment.

### IFN-signaling is required for Ly6C^hi^ monocyte maturation to FasL^+^MC phenotype

In order to determine the importance of IFN signaling for the maturation of Ly6C^hi^ monocytes into a FasL^+^MC phenotype, we infected *Stat2^-/-^* mice with H5N1 and examined surface marker expression. We found that Ly6C^hi^ monocytes present in *Stat2^-/-^* mice fail to acquire a FasL^+^MC phenotype – they express lower levels of Ly6C, do not upregulate CD11c, and express lower levels of MHCII. Paradoxically, despite being defective in mounting an IFN-dependent response, they also express lower levels of influenza M2 protein. Importantly they also fail to upregulate FasL expression (Fig 7A). Because H5N1-infected *Stat2^-/-^* mice fail to induce CCL2 and CCL7 and show only low levels of monocyte recruitment (35), it is unclear whether this represents the same population of cells present in WT mice. In order to determine if *Stat2^-/-^* monocytes could mature in the presence of chemokines, we examined the phenotype of these cells in mixed bone marrow chimeras. Although *Stat2^-/-^* Ly6C^hi^ monocytes could be recruited to the lung following H5N1 infection, they did not acquire a FasL^+^MC phenotype – they expressed lower levels of Ly6C and CD11c, lower levels of influenza M2, and failed to upregulate FasL (Fig 7B). Importantly no difference in expression of these markers was present in the absence of infection in Ly6C^hi^ monocytes, or in the absence or presence of infection in neutrophils. Ly6C^hi^ monocytes differentiate into Ly6C^lo^ monocytes, which are involved in tissue repair (10–12). Importantly, while *Stat2^-/-^* monocytes did not acquire an activated FasL^+^MC phenotype, they accounted for the majority of Ly6C^lo^ monocytes by day 3 post-infection in mixed bone marrow chimeras (Fig 6C and 7C). The data suggest that upon recruitment to the lungs, monocytes acquire a FasL^+^MC phenotype upon exposure to IFN, or differentiate into Ly6C^lo^ monocytes in the absence of IFN signaling.

**Figure 7.**
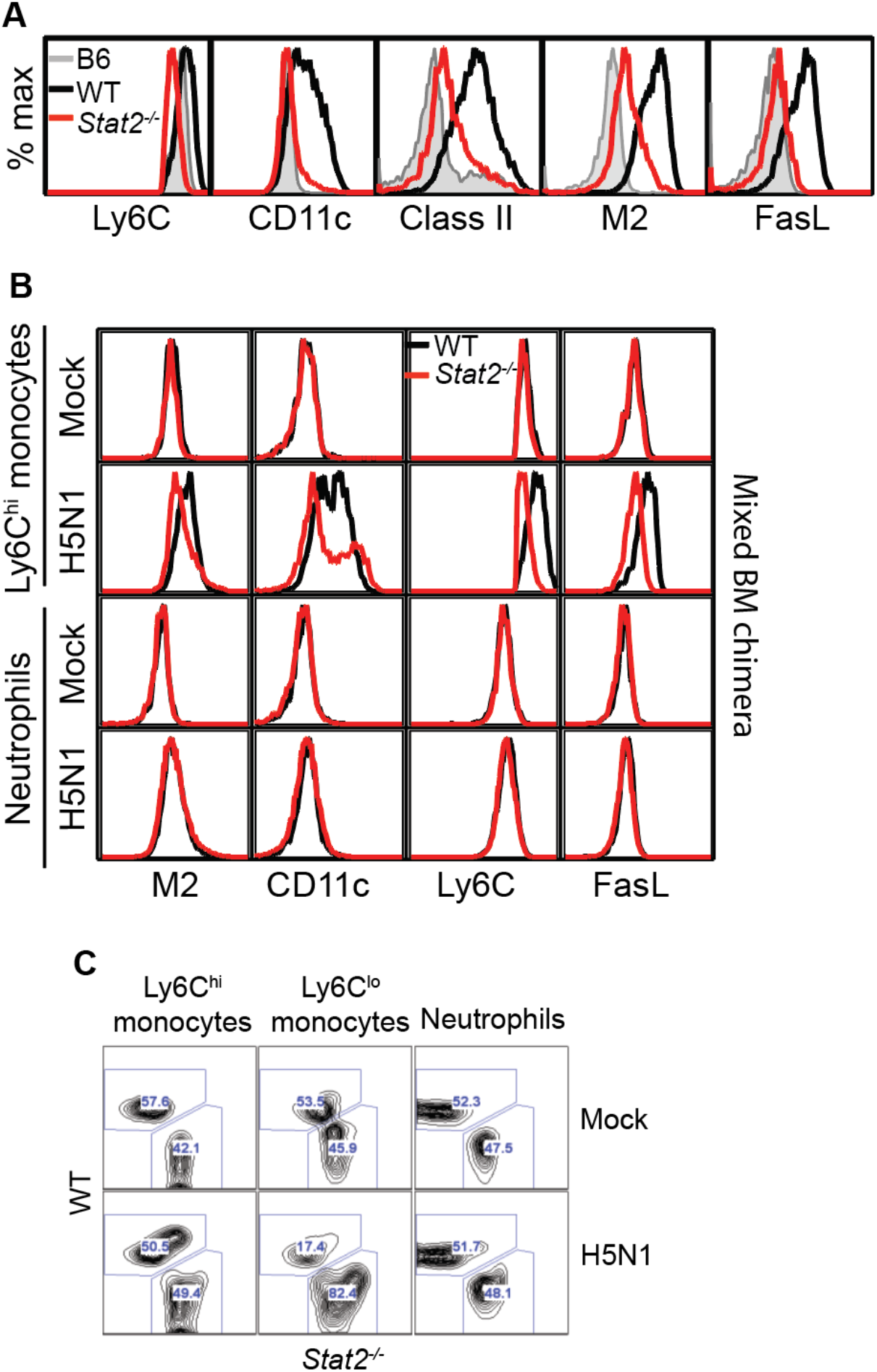
IFN signaling is required for maturation to FasL^+^MCs. **(A)** WT and *Stat2^-/-^* mice were infected with 10^2^ pfu of H5N1 and expression of the indicated markers on Ly6C^hi^ monocytes was determined at day 2 post-infection (n=3). **(B)** WT:*Stat2^-/-^→WT* 50:50 mixed bone marrow chimeras were infected with 10^2^ pfu of H5N1 and expression of the indicated markers on Ly6C^hi^ monocytes and neutrophils was determined by flow cytometry (n=4). Data are representative of 3 experiments for A and 2 for B.

## Discussion

### H5N1 infection is associated with monocytes, monocyte infection, and chemokines

H5N1 infection in humans is characterized by fever, respiratory symptoms, leukopenia, and rapid progression to pneumonia, followed by acute respiratory distress syndrome (ARDS) and multiple organ dysfunction. Lung pathology is characterized by extensive infiltration, diffuse alveolar damage and hyaline membrane formation characteristic of ARDS. The virus has been reported to disseminate systemically in some cases, however most patients die of respiratory failure. A distinctive feature of H5N1 infection is high serum cytokine and chemokine levels referred to as “cytokine storm,” which has been suggested to contribute to the severity of disease (42, 43). Consistent with chemokine levels, infiltrate primarily composed of monocytes, macrophages, and neutrophils is observed in the lung (49–51). The severe infection, systemic spread, and cytokine phenotypes are also recapitulated in mouse models of H5N1 (52). Here we show that very early following H5N1 infection high numbers of Ly6C^hi^ monocytes are found in the lungs. Many previous studies have identified infiltrating cells as macrophages by either histology or flow cytometry, but given the extent of influx and overlapping marker expression, these cells are likely MCs (22, 36, 37, 53). In addition, tissue-resident alveolar macrophage populations are rapidly depleted during influenza infection (54). Excessive monocyte infiltration has also been reported in SARS-CoV-2 (17–19). CCL2 and CCL7 production in our study correlates with the influx of Ly6C^hi^ monocytes, and previous studies have identified Ly6C^hi^ monocytes as the primary producers of these chemokines (22, 36). Human histological sections also confirm monocyte/macrophage chemokine expression (55). Studies have also linked CCL2 and CCL7 production to infiltrating monocytes during SARS-CoV-1 (17) and SARS-CoV-2 infection (56). In addition, we find extensive virus infection of Ly6C^hi^ monocytes during H5N1 infection. Virus infection has been observed during H5N1 infection primarily in epithelial cells and monocytes/macrophages (49, 55) and in monocytes during H7N9 infection (57), and productive macrophage replication has been suggested to be a unique feature of these viruses (58). We and others (22, 44) have observed monocyte infection with H1N1 strains, however in our experiments infection occurs at later time points in lower numbers of cells compared to H5N1 infection. Infection of macrophage subsets has also been reported in SARS-CoV-2 infection (https://doi.org/10.1101/2020.03.27.20045427). Whether infection and/or productive replication in monocytes are important determinants of H5N1 pathogenesis requires further investigation. Importantly, we also show that monocytes are the primary IFN producing and responding cells during infection; a finding also supported in vaccinia virus infection and lupus models (59, 60).

### Monocytes, FasL, TRAIL contribute to pathogenesis

We show that monocytes contribute to pathogenesis during H5N1 infection, as *Ccr2^-/-^* mice are protected from mortality. Although *Ccr2^-/-^* mice show delayed viral clearance and decreased T cell priming (20, 21), they are protected from excessive lung damage and lethal influenza infection in most studies (20, 22, 23). Treatment with CCR2 depleting antibody also protects from SARS-CoV-1 lethality (17). TRAIL expression by MCs referred to as exudate macrophages contributes to lethal influenza virus infection (23). In addition, higher IFN levels in some mouse strains leads to Ly6C^hi^ monocyte influx and TRAIL expression (26, 38). TRAIL neutralization protects from lung damage and death in both cases. Here we found FasL rather than TRAIL expression on Ly6C^hi^ monocytes. Expression of FasL during influenza virus infection has also been reported in other studies, and neutralization or genetic mutation protects from lethal infection (45, 46). We show MC expression of FasL, however FasL is also known to regulate T cell apoptosis during influenza virus infection (61), therefore it is possible that the effect of FasL on survival is mediated by cells other than monocytes. However, in the case of TRAIL-mediated tissue damage, T cell deficiency did not affect disease outcome (26). The reasons for differential expression of TRAIL and FasL are unknown but may be due variation in mouse or viral strains. Interestingly, neutralization of TRAIL or FasL during *in vitro* replication of influenza virus in epithelial cells leads to decreased viral titers, implying that the virus modulates cell death pathways to enhance replication (62). During SARS-CoV-1 infection, Ly6C^hi^ monocytes also express FasL in an IFN-dependent manner (17), however TNF-α rather than FasL contributes to pathogenesis in this case, highlighting that MCs can mediate pathogenic effects through various mechanisms. Importantly, monocytes are not universally pathogenic; they are protective during RSV, HSV, and WNV infection (48, 63).

### IFN not required for recruitment, differentiation to Ly6C^lo^

A number of studies have reported loss of Ly6C^hi^ monocyte recruitment in *Ifnar1^-/-^* mice following infection with influenza (22, 35, 36), SARS-CoV-1 (17), and SARS-CoV-2 (19), however we show that IFN signaling in monocytes is not directly required for recruitment. In contrast, another report showed lower levels of Ly6C^hi^ and higher levels of Ly6C^lo^ monocytes in WT: *Ifnar1^-/-^* mixed bone marrow chimeras (64). However, monocyte recruitment was examined at day 7 in this study, whereas we found approximately equal ratios of WT and *Stat2^-/-^* Ly6C^hi^ monocytes in mixed bone marrow chimeras at both day 2 and 3 post-infection. Given that we found an absence of IFN-signaling was associated with increased Ly6C^lo^ monocytes at day 3, it is likely that many of the recruited *Ifnar1^-/-^* Ly6C^hi^ cells had converted to Ly6C^lo^ cells by day 7. A number of labeling studies provide evidence that Ly6C^hi^ monocytes can give rise to Ly6C^lo^ monocytes (reviewed in (65)). However Ly6C^hi^ but not Ly6C^lo^ monocyte production is effected by the absence of the transcription factors IRF8 and KLF4 (8, 9), suggesting that alternate developmental pathways may exist. In addition, mice deficient for the transcription factor Nur77 lack Ly6C^lo^ monocytes specifically (66). The relative importance of these proposed developmental pathways is unknown and is likely to vary under pathologic conditions. In a mouse model of pristane-induced inflammation, IFN-signaling stimulated the production of chemokines that recruited Ly6C^hi^ monocytes to the peritoneum via CCR2. *Ifnar^-/-^* mice had high levels of Ly6C^lo^ monocytes consistent with our findings; labeling experiments supported a differentiation of Ly6C^lo^ cells from Ly6C^hi^ precursors (67). Regardless of whether Ly6C^lo^ monocytes differentiate from Ly6C^hi^ precursors, or develop independently, type I IFN signaling appears to be important for controlling the balance between the subsets. Ly6C^lo^ monocytes are generally thought to differentiate into resident macrophages that promote wound healing and resolution of inflammation. They produce only low levels of proinflammatory cytokines, and higher levels of anti-inflammatory factors including IL-1RA, IL10R, and ApoA/E, and CXCL16 (68). Ly6C^lo^ monocytes patrol the vasculature and remove damaged cells (68, 69). They also cross-present antigens derived from apoptotic cells, and promote tolerogenic responses through expression of PDL1 (70). How these cells contribute to influenza virus infection remains to be investigated.

### IFN required for monocyte maturation to MCs

*Stat2^-/-^* Ly6C^hi^ monocytes fail to differentiate into FasL^+^MCs, indicating that IFN signaling regulates the maturation of Ly6C^hi^ monocytes into MCs. Consistent with this, monocytes in human blood have been reported to be infected with a number of viruses, and differentiate into MCs that express dendritic cell markers in association with type I IFN production and ISG upregulation (17, 71, 72). IFN treatment has also been reported to lead to maturation of human monocytes into TRAIL^+^MCs, which induce IL-15 production and promote T cell responses (73). The Aryl hydrocarbon receptor (AhR) and IRF4 have been reported to be involved in differentiation of Ly6C^hi^ monocytes into MCs in both human and mouse experiments (74). Interestingly, we observed much lower levels of Ly6C^hi^ monocyte infection in *Stat2^-/-^* cells, implying that the differentiation to a MC phenotype in some way influences susceptibility to infection, an observation that is currently under investigation. Important roles for Ly6C^hi^ monocyte IFN signaling have also been defined in other infection models. Ly6C^hi^ monocytes control CD8+T cell and NK cell immunity through IFN-induced IL-18 and IL-15 production in response to a variety of bacterial, viral, and parasite infections (75). During HSV-2 infection, IFN signaling in inflammatory monocytes leads to IL-18 production, which subsequently activates NK cells that are critical for clearance of infection (76). In a *Candida* infection model, type I IFN-dependent IL-15 production by inflammatory monocytes in the spleen is critical for the activation of NK cells and neutrophils (77). Thus while IFN appears to have important effects on the maturation of Ly6C^hi^ monocytes into MCs, the resulting effects are protective in many cases, but pathogenic in some cases.

### Model

Altogether the data suggest a model for how type I IFNs contribute to H5N1 pathogenesis (Fig 8). Upon initial infection, influenza virus is detected by both epithelial and hematopoietic cells resulting in an initial wave of IFN-α/β and λ production. pDCs and alveolar macrophages have been defined as the primary IFN-producing cells at early time points following respiratory virus infection (78). This IFN is detected by resident Ly6C^hi^ monocytes, which signal through IFNAR1/2 to induce the production of ISGs including CCL2 and CCL7 – leading to an influx of Ly6C^hi^ monocytes from the bone marrow to the blood. Additional factors control the recruitment from the blood to the lung. Recruitment of these monocytes is not directly dependent on IFN, but is dependent on IFN-induced chemokine production. Once recruited to the lung Ly6C^hi^ monocytes mature into various MC effector populations dependent on the level of IFN. In the absence of IFN-signaling *Stat2^-/-^* cells become Ly6C^lo^, a phenotype associated with tissue repair. Intermediate levels of IFN likely facilitate beneficial outcomes including T cell priming and viral clearance. In the case of H5N1 infection, Ly6C^hi^ monocytes become infected, produce high levels of IFN-β and CCL2/7 – leading to excess influx, and mature into MCs that expresses CD11c, MHC Class II, and FasL facilitating tissue destruction.

**Figure 8.**
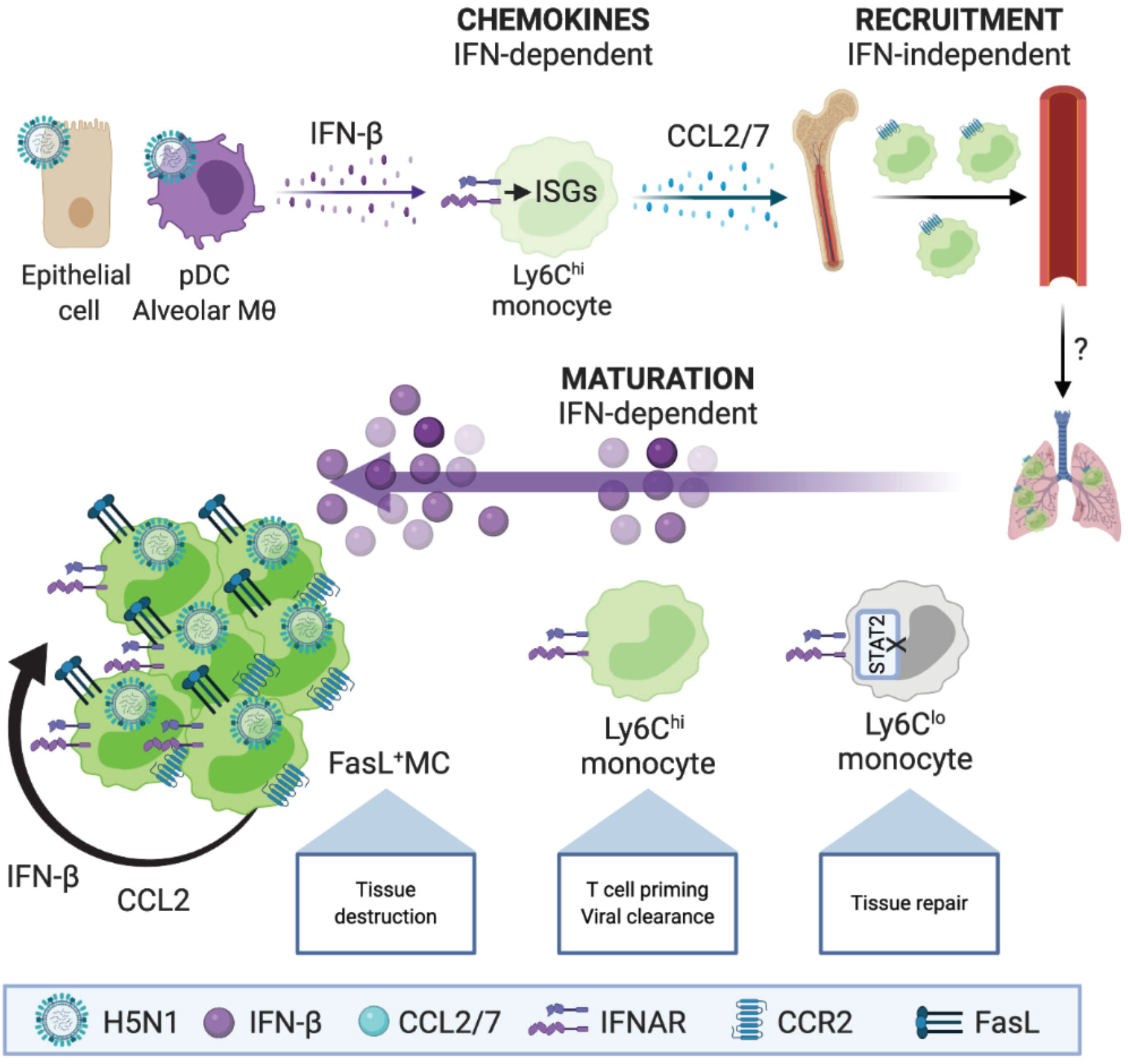
Model of the role of IFN in influenza virus infection. Following initial infection a primary wave of IFN is produced by epithelial cells, pDCs and alveolar macrophages. This induces the production of ISGs including CCL2 and CCL7, leading to Ly6C^hi^ monocyte recruitment to the lung – in a manner not directly dependent on IFN. Once recruited to lung, maturation of monocytes is regulated by the level of IFN. Lack of IFN-signaling in *Stat2^-/-^* cells leads to skewing to a Ly6C^lo^ phenotype, associated with tissue repair. Intermediate levels of IFN likely lead to T cell priming and viral clearance. During H5N1 infection, Ly6C^hi^ monocytes become infected, secrete high levels of IFN and chemokines – leading to further recruitment, and mature into a CD11c+MHCII+FasL+ MC phenotype and tissue destruction.

---

**Figure S1.**
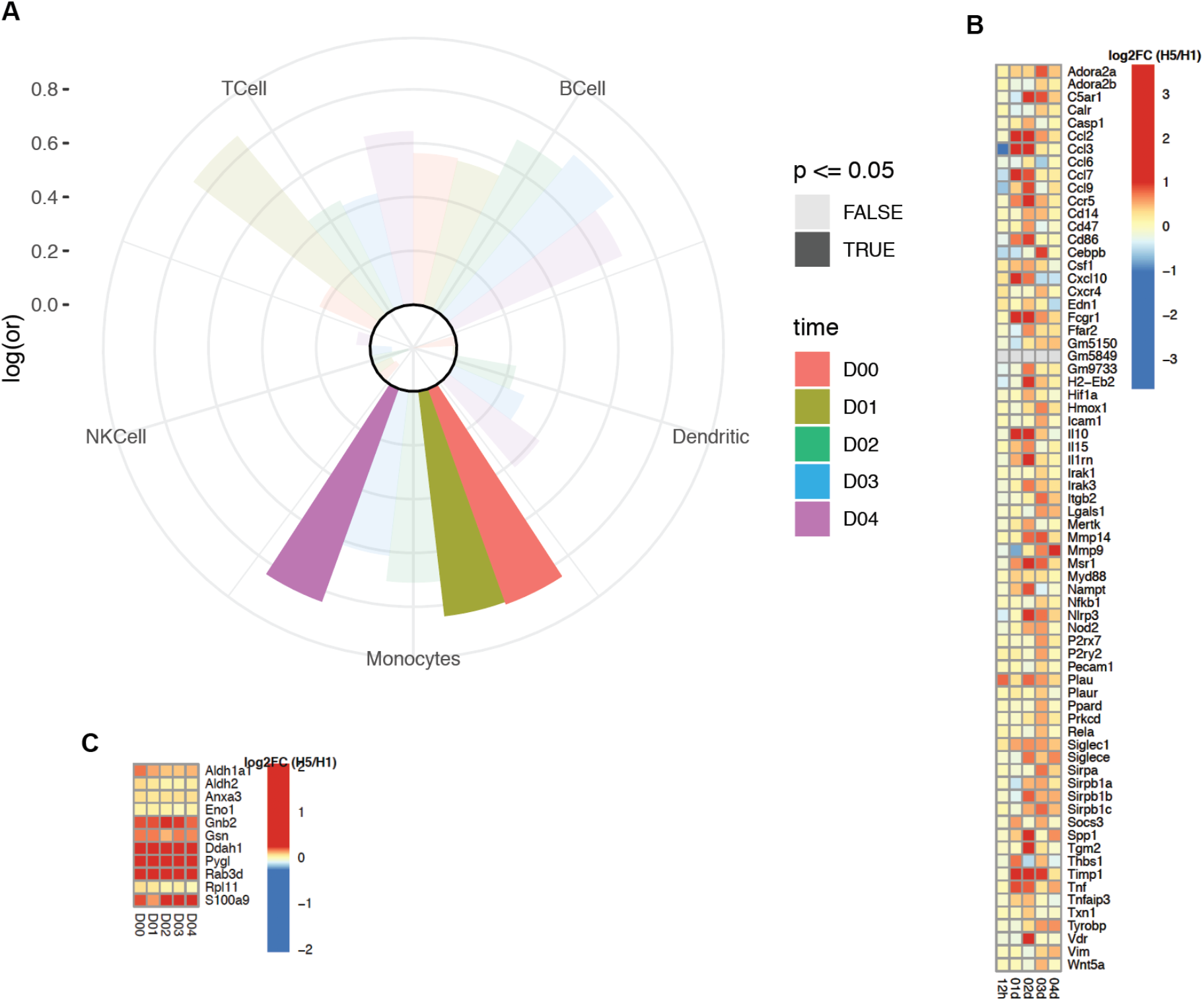
Proteic markers of monocytes are preferentially induced following H5N1 infection. (**A**) Fisher’s exact test was used to assess statistically the overlap between protein markers of immune cells (B, T, NK, Monocytes, DC) among proteins differentially expressed between H5N1- and H1N1-infected mice in lung tissue. The radial plot presents the log odd ratio (log(or)) of each set of markers (quadrants) for different timepoints investigated after infection (bars). A log odd ratio > 0 correspond to overlap of cell markers among proteins induced in H5N1 compared to H1N1 while a log odd ratio < 0 corresponds to the enrichment of cell markers among proteins repressed in H5N1 compared to H1N1. Benjamini-Hochberg correction was used to adjust for false positives. (**B**) Heatmap showing the log2 fold-change (log2FC) of monocytes transcriptomic markers between H5N1- and H1N1-infected mice in lung tissue. A blue-white-red color gradient depicts the gene the most repressed to the gene the most induced in H5N1-infected mice compared to H1N1-infected mice. (**C**) Heatmap showing the log2 fold-change (log2FC) of monocytes proteomic markers between H5N1- and H1N1-infected mice in lung tissue. A blue-white-red color gradient depicts the protein the most repressed to the protein the most induced in H5N1-infected mice compared to H1N1-infected mice.

**Figure S3.**
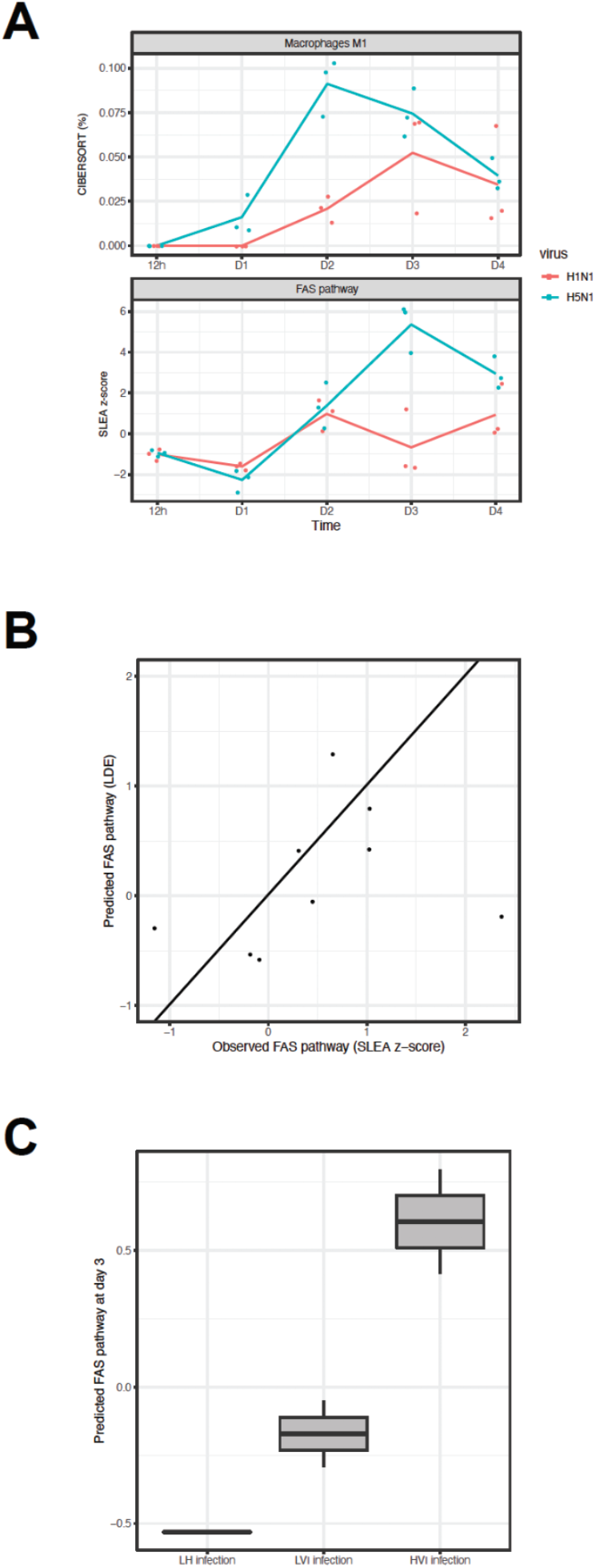
FAS pathway activation by M1 macrophages. (**A**) Inferred frequency of M1 macrophages (upper panel) and FAS pathway activation (lower panel) as a function of time. M1 macrophages recruitment was maximal two days following infection with H5N1 while FAS pathway was maximum activation was 1 day later (day three following infection with H5N1). (**B**) Mathematical modeling of FAS pathway at day d, was expressed as a linear function of FAS pathway activation at day d-1 and M1 macrophage frequency at day-1. The resulting model was confirmed on a publicly available set of transcriptomic data of mice infected with mock (LH), low virulent H3N2 (LVI) and highly virulent H3N2 virus (HVI). The activation of FAS pathway predicted by the mathematical model (y-axis) is presented as a function of the observed activation of FAS pathway (x-axis). The resulting Spearman correlation was 0.617 and a t-test p-value of 0.0857. (**C**) Boxplot of the activation of FAS pathway predicted by the mathematical model as a function of H3N2 virulence.

**Figure S2.**
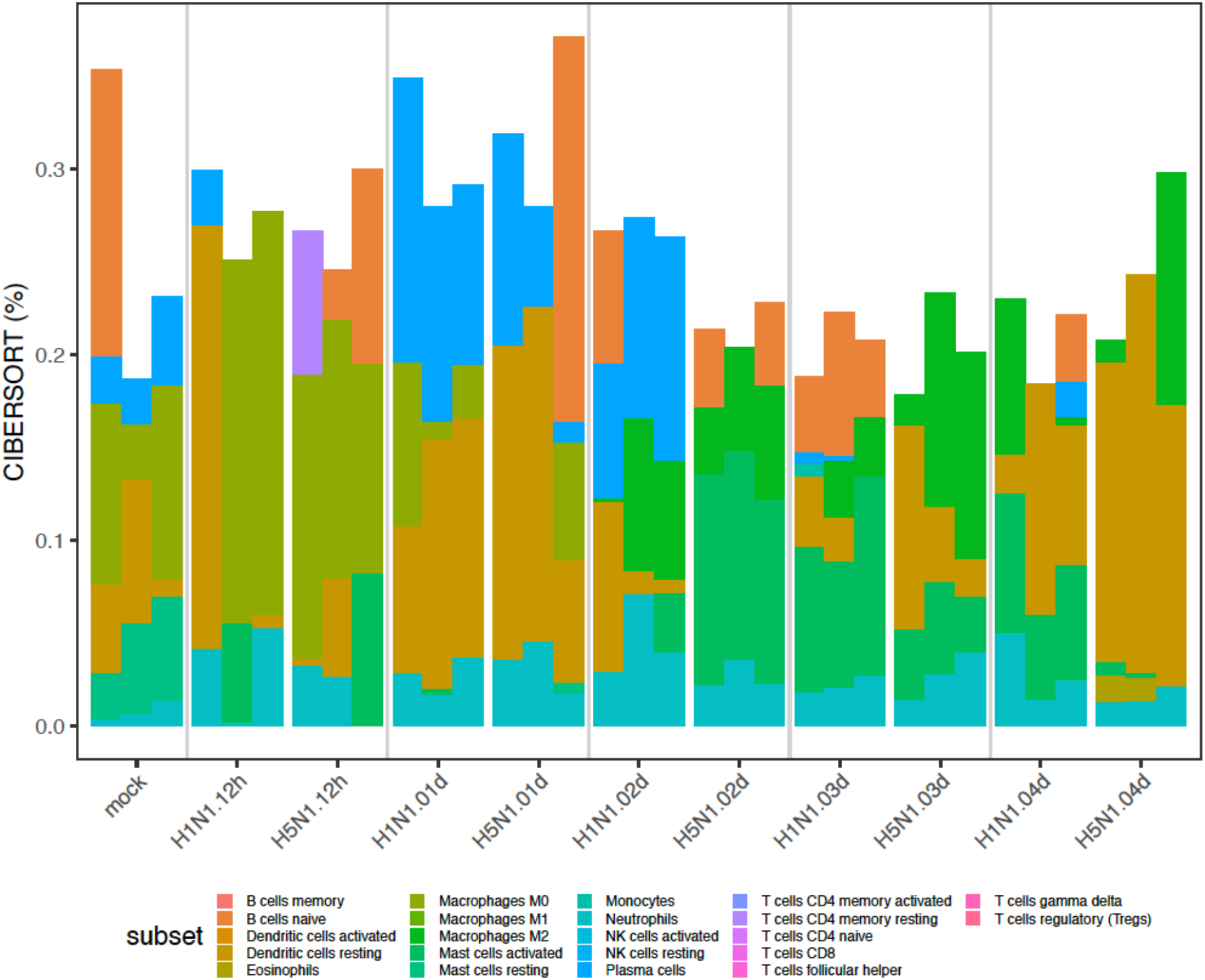
Cell subset frequencies in mice lung following influenza infections. Barplot of the relative frequencies of cell subsets inferred using CIBERSORT on the transcriptomic data collected after infection with H1N1, H5N1 or mock. The height of each bar is proportional with the inferred frequency of a subsets (color) in each triplicated samples (bars).

## Materials and Methods

### Mice

Animal studies were approved by the Institutional Animal Care and Use Committee of Icahn School of Medicine at Mount Sinai. C57BL/6J *Ifnar1^-/-^* (24), *Stat2^-/-^* (84), and *Mx1^gfp^* (paper 1) mice have been previously described. *Ifnb*^mob^, *Ccr2^-/-^*, and *Fasl^gld^* mice on the C57BL/6J background were purchased from Jackson. For mixed bone marrow chimeras 6-week old B6.SJL-Ptprc^a^ Pepc^b^/BoyJ (CD45.1) female mice were irradiated with 2 doses of 600 rads and injected with a 50:50 mix of WT (CD45.1) and *Stat2*-/- (CD45.2) bone marrow.

### Lung isolations and flow cytometry

Lungs were digested for 40 min in 1 mg/ml collagenase type 4 (Worthington) 5% FBS in DMEM. Cells were then filtered through a 0.2 μm cell strainer and RBCs were lysed. Cells were suspended in 3% FBS 2 mM EDTA in PBS and staining was performed in the presence of 2% NRS, 2% Fc block (BD), and fixable viability dye eFluor 450 (ebiosciences). Cells were stained with the following antibodies from BD: Ly6C-PerCP-Cy5.5 (AL-21), CD11b-PE (M1/70), Ly6G-V450 (1A8), CD11c-V450 or PE-Cy7 (HL3), CD45.1-FITC (A20), CD45.2-PE-CF594 (104), the following antibodies from eBiosciences: MHC Class II (I-A/I-E) e450 (M5/114.15.2), FasL-APC (MFL3), and from R&D: CCR2-APC (475301). Influenza M2 (E10) (85) was conjugated to Alexa 647. Cells were fixed with 2% formaldehyde after staining and analyzed on an LSRII after gating for FSC/SSC, singlets, and live cells.

### Infections and chemokine treatment

Mice were infected with the following viruses at the doses indicated in 20 μl PBS: A/PR/8/34 (H1N1) (PR8), A/Viet Nam/1203/04 (H5N1) lacking the multibasic cleavage site (HALo)(39). Viral titer was determined by plaque assay on MDCK cells. Mice were injected IV with 500 ng of CCL7 (47) and blood was harvested 30 min later in EDTA, RBCs were lysed and cells were analyzed by flow cytometry.

### qRT-PCR

Total RNA was extracted from collagenase-digested lung using using EZNA total RNA kit and RNase-free DNase (Omega). RNA was reverse-transcribed using Maxima Reverse Transcriptase and oligo-dT (Thermo). Quantitative RT-PCR was performed on cDNA using LightCycler 480 SYBR Green I Master Mix (Roche) and the primers ttgacccgtaaatctgaagctaat-*Ccl2F*, tcacagtccgagtcacactagttcac-*Ccl2R*, ggatctctgccacgcttctg-*Ccl7F*, tccttctgtagctcttgagattcctc-*Ccl7R*, cacagccctctccatcaacta-*Ifnb1F*, catttccgaatgttcgtcct-*Ifnb1R*, gtaacccgttgaaccccatt-*18SF*, and ccatccaatcggtagtagcg-*18SR* on a LightCycler 480 II and expressed as 2^-ΔΔCt^ relative to 18S.

### RNA-Seq analysis

Mouse lungs were homogenized in liquid nitrogen. 1 ml Trizol was added per 50-100 mg of tissue and the homogenate was incubated at room temperature for 5 minutes. 200 μl chloroform were added per ml of Trizol, shaken vigorously for 15-30 seconds and incubated at room temperature for 15 minutes. Samples were centrifuged for 15 minutes at 4°C and 12,000 rcf. The aqueous phase was carefully collected in a separate tube. 500 μl isopropanol was added per 1 ml Trizol, inverted to mix and incubated at room temperature for 5-10 minutes. Samples were transferred to Qiagen RNeasy Spin Columns and RNA was isolated according to the manufacturer’s instructions.

Libraries were constructed using the Illumina TruSeq Stranded Total RNA Library Prep Kit. Pair-end sequencing of 126 base pair reads was used. Raw reads were trimmed using Trimmomatic (version 0.36) using default value for pair-end sequencing. The trimmed reads were then aligned to the mouse genome (build GRCm38.91) using the STAR aligner (version 2.5.3a). Aligned reads were then counted using HTSeq (version 0.9.1). Differential expressed genes between H5N1- and H1N1-infected mice was determined by fitting a generalized linear model with gene counts as dependent variable and the virus strain (H5N1 or H1N1) as independent variable. A likelihood ratio test and Benjamini-Hoshberg correction, as implemented in the R package edgeR, was used to assess the statistical significance of the differential expression.

Gene Set Enrichment Analysis (GSEA) was used to assess the significance of MSigDB (version 6.2), blood-cell markers (PMID: 21743478, 28263321) and macrophages signatures (PMID: 28694385, 28421818), cytokines/chemokines (https://www.immport.org/resources/cytokineRegistry) and interferome (version 2) genesets. Gene were ordered from the most induced after H5N1 infection to the gene the most repressed by H5N1 infection compared to H1N1 using edgeR p-values (i.e. -log10(p) x sign(logFC)). Default parameters were used except the maximum genesets size was increased to 2000 genes and fixed seed for the permutation test equal 101. Cibersort (version 1.03) was used to infer cell subsets frequency based on RNA-Seq data.

CIBERSORT (version 1.03) was used to infer immune cell subset frequencies using default parameters.

### Microarray analysis

The human blood dataset of healthy individuals challenged with H1N1 was previously described in [PMID: 30356117]. Briefly, two independent microarray datasets (DEE3 H1N1, DEE4X H1N1) were combined. Each of these studies are from human viral exposure trials where healthy volunteers were followed for 7-9 days following controlled nasal exposure to H1N1 virus. Subjects enrolled into these viral exposure experiments had to meet several inclusion and exclusion criteria. Among them was an evaluation of pre-existing neutralizing antibodies to the viral strain. Any subject with pre-existing antibodies to the viral strain was excluded. All subjects exposed to H1N1 influenza received oseltamivir 5 days post-exposure. All subjects provided written consents, and each of the seven trials was reviewed and approved by the appropriate governing IRB. Symptom data and nasal lavage samples were collected from each subject on a repeated basis over the course of 7-9 days. Viral infection was quantified by measuring release of viral particles from nasal passages (viral shedding), as assessed from nasal lavage samples via qualitative viral culture and/or quantitative influenza RT-PCR. Symptom data were collected through self-report on a repeated basis. Symptoms were quantified using a modified Jackson score, which assessed the severity of eight upper respiratory symptoms (runny nose, cough, headache, malaise, myalgia, sneeze, sore throat, and stuffy nose) rated 0-4, with 4 being most severe. Scores were integrated daily over 5-day windows. Blood was collected and gene expression of peripheral blood was performed 1 day (24-30 h) prior to exposure, immediately prior to exposure, and at regular intervals following exposure. These peripheral blood samples were gene expression profiled on the Affy Human Genome U133A 2.0 array. Only raw (CEL files) gene expression data that pass QC metrics including those for RNA degradation, scale factors, percent genes present, β-actin 3’ to 5’ ratio and GAPDH 3’ to 5’ ratio in the Affy Bioconductor package were used for downstream analysis. Normalization via RMA was performed on all expression data across all timepoints. The primary endpoint used for differential expression analysis was the symptom score as continuous variable indicating the log of the maximum 5-day integrated symptom score +1. Linear regression using the R package LIMMA was used to identify genes correlated to the symptom score. GSEA on the gene list ranked by the LIMMA t-static and using the blood-cell markers (PMID: 21743478, 28263321) as reference database was used to identify immune cell subsets correlated to the symptom score.

### Proteomics

Protein abundance in the lung homogenates was measured using amine-reactive isobaric tags in an untargeted run. Briefly, after lysis and protein digestion, peptides from the lung homogenates were labeled with a Tandem Mass Tag (TMT; Thermo Fisher) and mixed in equal proportions. These mixed samples were analyzed by LC-MS on a Thermo Orbitrap Fusion with extended run times to ensure deep coverage. All samples were prepared in biological triplicate and run in technical duplicate to increase sequencing depth. Peptide searches using MaxQuant and statistical testing using MSstatTMT were performed. Fisher exact test was used to assess enrichment of cell subset markers among proteins differentially expressed between H5N1- and H1N1-infected mice.

### Statistical analysis

A Wilcoxon rank-sum test was used to compare levels between two groups. A Spearman correlation and a Student t-test was used to assess the correlation between two continuous variables.

## Author contributions

MBU conceived the project, performed the experiments, generated the figures and wrote the manuscript. AGS supervised the work, provided funding, and critically reviewed the manuscript.

## Acknowledgements

We thank Christian Schindler for originally providing the *Stat2^-/-^* mice. We are grateful to Richard Cadagan and Osman Lizardo for excellent technical assistance. This work was partially supported by CRIP (Center for Research on Influenza Pathogenesis), an NIAID funded Center of Excellence for Influenza Research and Surveillance (CEIRS, contract # HHSN272201400008C to AG-S, RA and MBU).

## DATA AND SOFTWARE AVAILABILITY

### Data availability

RNA-Seq have been deposited in the NCBI’s Gene Expression Omnibus with the accession number GSE98527. Proteomics data have been deposited in MassIVE and will be made available with publication of the article.

### Code availability

All the source code used to generate the figures part of this manuscript is available at “https://github.com/sekalylab/fluomics.lung”. The authors declare that all other data supporting the findings of this study are available from the authors upon request.

